# *PhenoGMM*: Gaussian mixture modelling of microbial cytometry data enables efficient predictions of biodiversity

**DOI:** 10.1101/641464

**Authors:** Peter Rubbens, Ruben Props, Frederiek-Maarten Kerckhof, Nico Boon, Willem Waegeman

**Affiliations:** KERMIT, Department of Data Analysis and Mathematical Modelling, Ghent University, Coupure Links 653, B-9000, Gent, Belgium; Center for Microbial Ecology and Technology (CMET), Ghent University, Coupure Links 653, B-9000, Gent, Belgium; Flanders Marine Institute (VLIZ), InnovOcean site, Wandelaarkaai 7, 8400, Ostend, Belgium

## Abstract

Microbial flow cytometry allows to rapidly characterize microbial communities. Recent research has demonstrated a moderate to strong connection between the cytometric diversity and taxonomic diversity based on 16S rRNA gene amplicon sequencing data. This creates the opportunity to integrate both types of data to study and predict the microbial community diversity in an automated and efficient way. However, microbial flow cytometry data results in a number of unique challenges that need to be addressed. The results of our work are threefold: i) We expand current microbial cytometry fingerprinting approaches by proposing and validating a model-based fingerprinting approach based upon Gaussian Mixture Models, which we called *PhenoGMM*. ii) We show that microbial diversity can be rapidly estimated by *PhenoGMM*. In combination with a supervised machine learning model, diversity estimations based on 16S rRNA gene amplicon sequencing data can be predicted. iii) We evaluate our method extensively by using multiple datasets from different ecosystems and compare its predictive power with a generic binning fingerprinting approach that is commonly used in microbial flow cytometry. These results demonstrate the strong connection between the genetic make-up of a microbial community and its phenotypic properties as measured by flow cytometry. Our workflow facilitates the study of microbial diversity and community dynamics using flow cytometry in a fast and quantitative way.

**Importance:** Microorganisms are vital components in various ecoystems on Earth. In order to investigate the microbial diversity, researchers have largely relied on the analysis of 16S rRNA gene sequences from DNA. Flow cytometry has been proposed as an alternative technique to characterize microbial community diversity and dynamics. It is an optical technique, able to rapidly characterize a number of phenotypic properties of individual cells. So-called fingerprinting techniques are needed in order to describe microbial community diversity and dynamics based on flow cytometry data. In this work, we propose a more advanced fingerprinting strategy based on Gaussian Mixture Models. When samples have been analyzed by both flow cytometry and 16S rRNA gene amplicon sequencing, we show that supervised machine learning models can be used to find the relationship between the two types of data. We evaluate our workflow on datasets from different ecosystems, illustrating its general applicability for the analysisof microbial flow cytometry data. *PhenoGMM* facilitates the rapid characterization and predictive modelling of microbial diversity using flow cytometry.

## Introduction

Various tools have been developed to study and monitor microbial communities. With the emergence of 16S rRNA gene sequencing, researchers have uncovered the genotypic diversity of microbial communities to a large extent [58]. However, microorganisms with the same genotype can still present different phenotypes, displaying so-called phenotypic heterogeneity [1]. Therefore, instead of solely focusing on genotypic information, there is a need to combine omics data with phenotypic information [17]. One of such tools to study the phenotypic identity of microbial communities is flow cytometry (FCM). FCM is a high-throughput technique, measuring hundreds to thousands of individual cells in mere seconds. These measurements result in a multivariate description of each cell, derived from both scatter and fluorescence signals. The first is related to cell size and morphology, while the latter depends on either autofluorescence properties or the interaction between the cell and a specific stain. Common for microbial FCM is to use a stain that interacts with the nucleic acid content of a cell [30, 59]. The combination of both FCM and 16S rRNA gene amplicon sequencing has been increasing in recent microbial surveys, which allows to investigate functional properties such as bacterial production or activity [9, 10, 53] but also to remove the compositional nature of microbiome data [44, 60].

Many algorithms exist in the field of immunophenotyping cytometry to identify separated cell populations (see e.g. the extensive benchmark studies by [2] and [62]). However, microbial cytometry data has a number of different characteristics, which is why most of these approaches are not applicable. This originates from the fact that bacterial cells are typically much smaller in both cell size and volume compared to eukaryotic cells [49], which complicate their detection. In addition, no general antibody-based panels have been established for microbial cells due to the high complexity of microbial communities [32]. One has to rely on general DNA stains, for which it is difficult to develop multicolor approaches [13]. Therefore, the number of variables describing an individual bacterial cell is typically much lower than for example a human cell. As a result, cytometric distributions of bacterial populations can highly overlap [51], as the number of bacterial populations is much larger than the number of differentiating signals. Therefore, automated cell population identification algorithms cannot be directly applied for the analysis bacterial cytometry data. Consequently, data analysis pipelines should be designed to address these characteristics.

To do so, microbiologists commonly rely on so-called cytometric fingerprinting techniques [30, 45]. Such a fingerprint allows to derive community-level variables in terms of the number of bins or clusters (i.e. gates), cell counts per cluster and the position of those clusters [31], despite the fact that there are no or only a few clearly separated cell populations. The approaches that are currently used for the analysis of bacterial communities can be broadly divided in two categories: i) manual annotation of clusters [23, 30] and ii) automated approaches that employ binning strategies [20, 29, 34, 45]. Current methods have a number of drawbacks: i) Manual gating of regions of interest is laborious in time and operator dependent, ii) traditional binning approaches result in a large number of variables (e.g., a fixed grid of dimensions 100 *×* 100 will result in 10,000 sample-describing variables) and because of that iii) only bivariate interactions of cytometry channels are considered when employing a traditional binning approach.

After a fingerprint has been constructed, one can calculate what is called the *cytometric* or *phenotypic* diversity of a community [34, 45]. These are estimations of the diversity of a microbial community based on the cell counts per cluster. If many clusters contain cells, a community can be considered as ‘rich’. If the cells are equally distributed over those clusters, a community can be considered as ‘even’. Recent reports have shown a significant correlation between the cytometric diversity and genotypic diversity derived from 16S gene sequencing data [20, 45, 46]. In other words, there is a strong connection between the genetic make-up of a microbial community and its phenotypic properties, which can be quantified. This result has been backed up by molecular identification using DGGE of sorted subpopulations [30, 41], the sequencing of sorted individual cells or subpopulations [22, 56, 63] and by using a bottom-up approach in which individual bacterial populations resulted in unique cytometric characterizations, which can be automatically identified using supervised machine learning models [51].

We propose an extension of current fingerprinting approaches in order to deal with many highly overlapping cell populations. Our workflow, which is called ‘*PhenoGMM* ‘, makes use of Gaussian mixture models (GMM). GMMs have been successfully applied to cytometry data before to identify separated cell populations in an automated way [8, 47]. Interestingly, Hyrkas et al. have shown that their GMM approach outperformed state-of-the-art immunophenotyping cytometry algorithms for the automated identification of phytoplankton populations [28]. By overclustering the data, GMMs can also be used to describe the distribution of the data, and therefore *PhenoGMM* is able to deal with overlapping cell populations. In addition, the number of mixtures that are needed to describe the data is much lower compared to the number of variables that result from traditional binning approaches. This facilitates the use of supervised machine learning models.

The aims of this study are the following. i) We expand current microbial cytometry fingerprinting methodologies by proposing a fingerprinting strategy based upon Gaussian Mixture Models. ii) We demonstrate that bacterial diversity can be rapidly estimated based on the cytometric fingerprints derived from *PhenoGMM*. iii) Fingerprints can also be used to predict diversity values based on 16S rRNA gene amplicon sequencing in combination with a supervised machine learning model. iv) We evaluate its performance both for synthetic and natural freshwater communities, and compare its performance with the predictive power of a generic traditional binning approach, which we have called *PhenoGrid*. v) Finally, we highlight a number of possible extensions concerning the integration of FCM with 16S rRNA gene sequencing, such as the calculation of *β*-diversity values and the prediction of individual OTU abundances based on FCM data.

## Materials and methods

### Methodology

#### Preprocessing

Two steps are carried out for all measurements before further analysis of the data. First, all individual channels are transformed using *f* (*x*) = asinh(*x*). Next, background due to debris and noise is removed using a fixed digital gating strategy [43, 45]. In other words, a single gate is applied to separate bacterial cells from background and is used for all samples.

#### Cytometric fingerprinting using Gaussian Mixture Models

In order to create a fingerprint template that can be used to extract variables describing a specific sample, all samples in the dataset in the training set need to be concatenated. Files are first subsampled to the same number of cells per file (`N_CELLS MIN`), in order to not bias the Gaussian Mixture Model (GMM) towards a specific sample. This number can either be the lowest number of cells present in one sample, or a number of choice. A rough guideline can be to not let the training set be larger than 1 *×*10^6^ cells, depending on computational resources. If *n* denotes the total number of samples, then the total number of cells (`N CELLS`) in the training set will be determined as `N_CELLS` = *n ×* N_REPN CELLS MIN, in which `N_REP` denotes the number of technical replicates of a specific sample. Typically, forward (FSC) and side scatter (SSC) channels are included, along with one or two targeted fluorescence channels (denoted as FL*X*, in which *X* indicates the number of a specific fluorescence detector). Unless noted otherwise, channels FSC-H, SSC-H and FL1-H (488 nm) are included for data analysis.

Once this training set is created, a GMM of *K* mixtures is fitted to the data. If **X** denotes the entire data matrix or training set containing *N* cells, then **X** consists of cells written as ***x***_1_, *…*, ***x***_*N*_, of which each cell is described by *D* variables (i.e., the number of signals collected from the flow cytometer). Cell *i* is described as 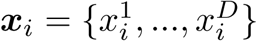. A GMM consists out of a superposition of normal distributions 𝒩, of which each distribution has its own mean *µ* and covariance matrix Σ. Each mixture has a mixing coefficient or weight *π*, which represents the fraction of data each mixture is describing. The distribution *p*, can be written as follows:

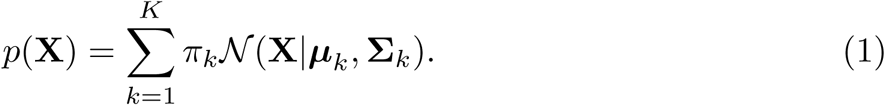

The set of parameters 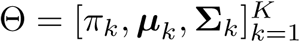 is determined by the expectation-maximization (EM) algorithm [7]. Once a GMM has been trained on the concatenated data, one can cluster cells per cluster and sample accordingly, by assigning each cell to the cluster for which it has the highest posterior probability. For this step, either a specific number of cells of choice are sampled per replicate, or the lowest number of cells of the replicates that are part of that specific sample, denoted as `N_CELLS_REP`. After clustering, we count the number of cells per cluster, after which the relative number of cells per cluster and sample can be retrieved. An illustration of *PhenoGMM* can be seen in Fig. 1.

**Fig 1.**
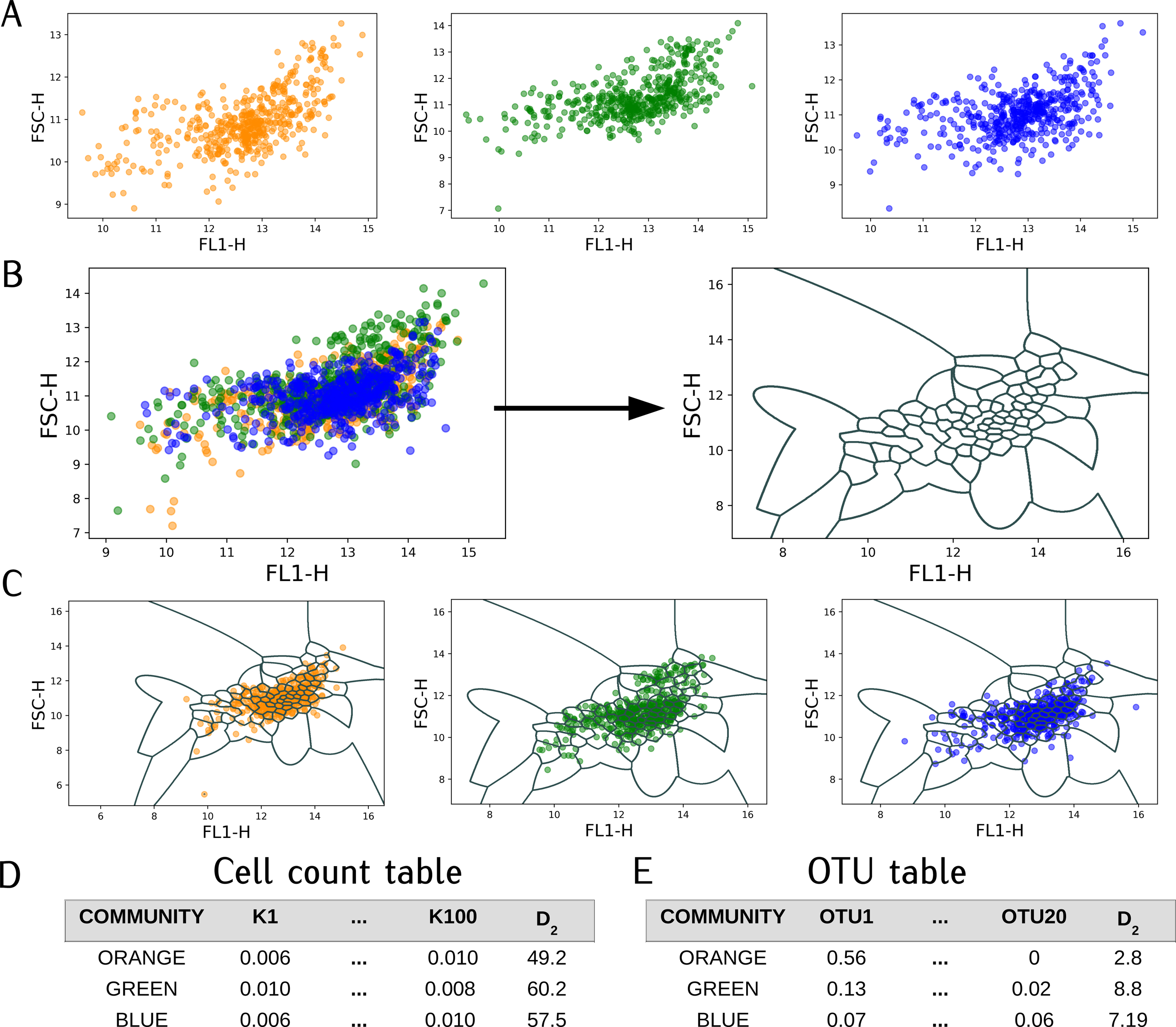
Illustration of *PhenoGMM* for two channels (FL1-H and FSC-H) using *K* = 100 mixtures. **A**: The analysis starts from cytometric measurements of three bacterial communities of interest, noted as ‘ORANGE’ (*S* = 6), ‘GREEN’ (*S* = 8) and ‘BLUE’ (*S* = 15). **B**: Data of the three communities are concatenated into one dataframe, to which is a GMM with *K* = 100 mixtures is fitted. This results in a cluster structure, which is depicted on the right. **C**: The fingerprint template is used to derive relative cell counts per cluster and per bacterial community. **D**: This results in a ‘count’ table, which can be used to rapidly quantify the cytometric diversity based on equations 2-4 (in this case *D*_2_). **E**: Based on the count table derived using *PhenoGMM*, one can try and predict diversity metrics based another type of data such as 16S rRNA gene sequencing, using a machine learning model.

We used the GAUSSIANMIXTURE() function of the scikit-learn machine learning library to implement our method [42]. This function contains four different ways in which the covariance matrix of each mixture is determined:

- `diag:` each mixture has its own diagonal covariance matrix.
- `full:` each mixture has its own general covariance matrix.
- `spherical:` each mixture has its own single variance.^1^
- `tied:` all mixtures share the same general covariance matrix.

Unless otherwise noted, we let each mixture have its own general covariance matrix (`full`). MCLUST was used to integrate *PhenoGMM* in the R package PhenoFlow [54].

#### Defining *α***- and** *β***-diversity**

Both 16S gene amplicon sequencing and flow cytometry fingerprints give rise to a compositional representation of a microbial community. The first is determined by counting the number of similar sequences at a certain taxonomic level, i.e. a taxonomic unit, the latter by counting the number of cells present in a predefined gate or cluster in the cytometric fingerprint, the phenotypic unit. Based on abundance data, one can calculate both *α*- and *β*-diversity metrics. The first quantifies the diversity within a sample, the latter the diversity between samples. Various diversity metrics exist in ecology to calculate *α*-diversity; we use the Hill numbers to quantify community diversity [26], as proposed by recent reviews [33] and [15]. If we let **p** = *p*_1_, *…, p*_*S*_ represent the vector of relative abundances, describing the abundance of *S* bacterial populations, then we can define the richness (*D*_0_) and evenness (*D*_1_, *D*_2_) of a microbial community as follows:

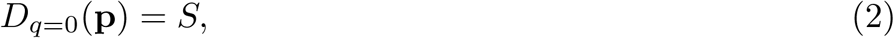

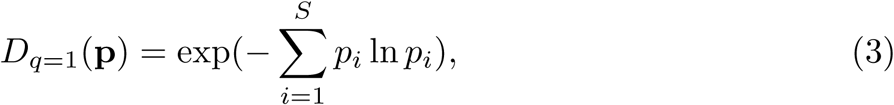

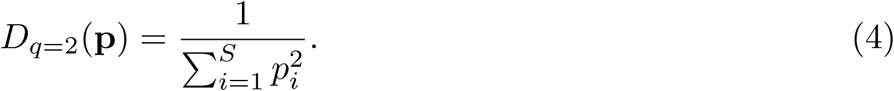

*q* denotes the order of the Hill-number, which is part of a general family, which can be denoted as 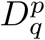. It expresses the weight that is given to more abundant populations.

*β*-diversity quantifies the difference in compositions between different samples. Typically, this is calculated by performing ordination on a dissimilarity matrix that contains the dissimilarities or distances between samples. We quantify the dissimilarity between samples using the Bray-Curtis dissimilarity [11]. If we let *BC*_*AB*_ denote the dissimilarity between samples *A* and *B, BC*_*AB*_ is calculated using the following equation:

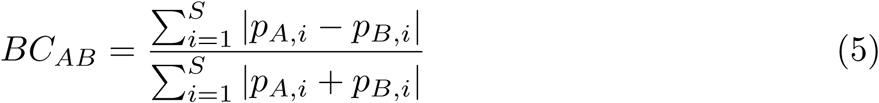

#### Predictive modeling

FCM fingerprints can be used as input variables to train a machine learning model. We use Random Forest regression [12], an ensemble of decision trees, to predict *α*-diversity metrics, based on an *in silico* community assembling strategy (see further) or 16S rRNA gene amplicon sequencing. A randomized grid search is performed to search for an optimal hyperparameter combination [5]. This means that a number of random combinations of hyperparameter values were evaluated. The maximum number of variables that are considered at an individual split for a decision tree is randomly drawn from {1, *…, K*}, the minimum number of samples for a specific leaf is randomly drawn between {1, *…*, 5}. The cross-validation strategy differs per experiment, and is described accordingly.

### Datasets

#### Dataset 1: In Silico Bacterial Communities

Data from 20 individual bacterial populations that were measured through FCM were collected from [51]. The data is available at FlowRepository (accession ID:

FR-FCM-ZZSH). In brief, bacterial populations were sampled after 24h of incubation, stained with SYBR Green I and two technical replicates per population were measured on an Accuri C6 (BD Biosciences). Fluorescence was measured by the targeted detector (FL1, 530/30 nm) and three additional detectors, next to forward (FSC) and side scatter (SSH) information that was collected as well. Additional automated denoising was performed using the FlowAI package (v1.4.4., default settings, target channel: FL1, changepoint detection: 150, [40]). A full experimental overview can be found in [51]. The lowest number of cells collected after background removal amounted to 13166 cells.

#### Dataset 2: Cooling water microbiome

Data was used as presented in [45]. Samples were collected from the cooling water of a discontinuously operated research nuclear reactor. This reactor underwent four phases: control, startup, operational and shutdown. Samples were taken from two surveys (Survey I and II) and analyzed through 16S rRNA gene amplicon sequencing (*n* = 77) and FCM (*n* = 153). The sequencing and flow cytometric procedures and data processing are extensively described in [45]. Taxonomic identification of the microbial communities was done at the operational taxonomic unit (OTU) level at 97% similarity. Sequences are available from the NCBI Sequence Read Archive (SRA) under (accession ID: SRP066190), flow cytometry data is available from FlowRepository (accession ID: FR-FCM-ZZNA). The lowest number of cells collected after background removal amounted to 10565 cells.

#### Dataset 3: Freshwater lake system microbiome

A total of 173 samples collected from three types of freshwater lake systems were analyzed. Data were used as presented in [53]. All samples were analyzed through 16S rRNA gene amplicon sequencing and FCM. Samples originate from three different freshwater lake systems: (1) 49 samples from Lake Michigan (2013 & 2015), (2) 62 samples from Muskegon Lake (2013-2015; one of Lake Michigan’s estuaries), and (3) 62 samples from twelve Inland lakes in Southeastern Michigan (2014-2015). Field sampling, DNA extraction, DNA sequencing and processing are described in [14]. Fastq files were submitted to NCBI SRA under BioProject accession number PRJNA412984 and PRJNA414423. Sequence data was processed based on the MiSeq standard operating procedure (see [53]). Taxonomic identification of microbial communities was done for each of the three lake datasets separately and treated with an OTU abundance threshold cutoff of either 1 sequence in 3% of the samples. For comparison of taxonomic abundances across samples, each the three datasets were then rarefied to an even sequencing depth, which was 4,491 sequences for Muskegon Lake samples, 5,724 sequences for the Lake Michigan samples, and 9,037 sequences for the Inland lake samples. Next, the relative abundance at the OTU level was calculated by taking the count value and dividing it by the sequencing depth of the sample. Flow cytometry procedures are extensively described in [46]. In brief, samples were stained with SYBR Green I and three technical replicates were measured on an Accuri C6 (BD Biosciences). Flow cytometry data is available from FlowRepository (accession IDs: FR-FCM-ZY9J and FR-FCM-ZYZN). The lowest number of cells collected after denoising amounted to 2342 cells.

#### Code & data availability

All code and data supporting this manuscript is freely available on GitHub at: https://github.com/prubbens/PhenoGMM. The functionality of *PhenoGMM* has been incorporated in the R package *PhenoFlow*: https://github.com/CMET-UGent/Phenoflow_package. Raw flow cytometry data is freely available on FlowRepository (accession numbers FR-FCM-ZZSH, FR-FCM-ZZNA, FR-FCM-ZY9J and FR-FCM-ZYZN). Raw sequences are avaialable via the NCBI Sequence Read Archive (accession numbers SRP066190, PRJNA412984 and PRJNA414423).

### Experimental setup

Our proposed fingerprinting approach based on GMMs was compared to a generic fixed binning approach, which we have called *PhenoGrid*. In brief, we implemented a binning grid of *L* = 128 *×* 128 for each bivariate parameter combination, after which cell fractions per bin were determined. The resulting cell fractions were next vectorized, concatenated and normalized. Both *PhenoGMM* and *PhenoGrid* result in multiple variables that describe relative cell counts, either per cluster or bin. We tested the following possibilities:

1. Unsupervised *α*-diversity estimations, by directly calculating *D*_0_, *D*_1_ and *D*_2_ according to equations 2, 3 and 4 based on the cell count vectors.
2. Unsupervised *β*-diversity estimations, by calculating Bray-Curtis dissimilarities (equation 5) between the cytometric fingerprints.
3. Supervised *α*-diversity predictions, with cytometric fingerprints as input variables to predict true target variables *D*_0_, *D*_1_ and *D*_2_ based on 16S rRNA gene sequencing data, by means of Random Forest regression.
4. Supervised taxon abundance predictions, with cytometric fingerprints as input variables to predict true taxon abundances, based on 16S rRNA gene sequencing data, by means of Random Forest regression.

#### *α*-diversity estimations of *in silico* synthetic microbial communities

In the first experiment, we assessed how well *PhenoGMM* was able to capture community variations in synthetic microbial communities. The main goal was to estimate (i.e., unsupervised) or predict (i.e., supervised) *α*-diversity metrics based on cytometric fingerprinting of the data. To do so, we performed an *in silico* community assembly strategy. In other words, cytometric characterizations of individual bacterial populations were artificially aggregated according to predefined compositions (conform [51]). The compositions were determined according to the following strategy:

1. Sample uniformly at random a number 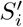 between two and 20; this is the number of populations that will make up community *i*.
2. Select randomly which 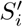 populations will make up the total community (from *S* = 20 populations).
3. Use the Dirichlet distribution to randomly sample a specific composition that sums to 1, containing the selected populations. The Dirichlet distribution can be used to model the joint distribution of individual fractions of multiple species [19]. The evenness of the composition depends on the concentration parameter *a*, which determines how evenly the weight will be spread over multiple species. If *a* is low, only a few species will make up a large part of the community. If *a* is high, the fraction of each population will be almost equally divided.

This was done using Dataset 1, which contains the cytometric characterization of individual bacterial populations. As it was known which cell belongs to which species, diversity indices could be calculated with high accuracy by simply counting the number of bacterial populations that were present in a community (*D*_0_) or by counting the fraction of cells that came from every population (*D*_1_, *D*_2_). We created a training set containing 300 different *in silico* communities and a test set containing 100 different communities. This experiment was repeated for *a* = 0.1, 1, 10. Random forests were trained using 5-fold cross-validation by means of a randomized grid search in which a 100 hyperparameter combinations are evaluated. Both unsupervised and supervised *α*-diversity estimations were reported for the test sets.

#### *α*-diversity estimations of natural microbial communities

Analogous to experiment 1, the goal was to both estimate and predict *α*-diversity metrics based on cytometric fingerprinting of the data. However, different from dataset 1, we now used *α*-diversity values based on 16S rRNA gene amplicon sequencing, as we considered natural microbial communities. Dataset 2 and 3 contain natural communities that were measured both by FCM and 16S rRNA gene amplicon sequencing. These values were used as target variables to predict. 10-fold blocked cross-validation was used to select hyperparameters for the Random Forest model [48], using a randomized grid search in which 50 combinations are evaluated. Samples were grouped in time (Dataset 2) or according to the location and the season (Dataset 3) in which they were measured. We reported the out-of-bag (OOB) predictions of the best Random Forest model. Unsupervised estimations were reported based on the full dataset.

#### Performance evaluation

- Unsupervised and supervised *α*-diversity estimations were quantified by calculating Kendall’s rank correlation coefficient *τ* between the true and estimated values. The *τ*_*B*_ implementation, which is able to deal with ties, was calculated as follows:

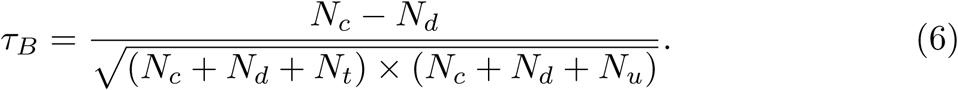

*N*_*c*_ denotes the number of concordant pairs between true and predicted values, *N*_*d*_ the number of discordant pairs, *N*_*t*_ the number of ties in the true values and *N*_*u*_ the number of ties in the predicted values. Values range from −1 (perfect negative association) to +1 (perfect positive association) and a value of 0 indicates the absence of an association. This was done using the KENDALLTAU() function in Scipy (v1.0.0).
- Supervised predictions were evaluated by calculating the *R*^2^ between true (**y** = *{y*_1_, *…, y*_*n*_*}*) and predicted (**ŷ** = *{ŷ*_1_, *…, ŷ*_*n*_*}*) values:

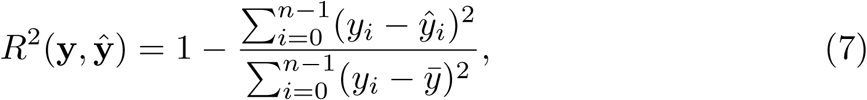

in which 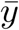 denotes the average value of **y**. If *R*^2^ = 1, predictions are correctly estimated. If *R*^2^ *<* 0, predictions are worse than random guessing.The R2 SCORE()-function from the scikit-learn machine learning library was used.
- Unsupervised *β*-diversity estimations were evaluated by calculating the correlation between Bray-Curtis dissimilarity matrices (*BC*) based on FCM and 16S rRNA gene sequencing data using a Mantel-test [37]. This test assesses the alternative hypothesis that the distances between samples based on cytometry data are linearly correlated with those based on 16S rRNA gene sequencing data. It makes use of the cross-product term *Z*_*M*_ across the two matrices for each element *ij*:

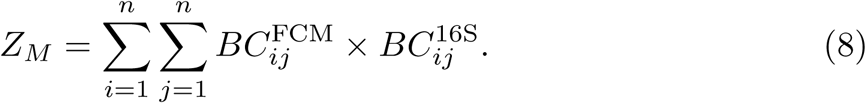 The test statistic *Z*_*M*_ is normalized and then compared to a null distribution, based on 1000 permutations.

## Results

### *PhenoGMM* allows to predict *α*-diversity of *in silico* synthetic microbial communities

In the first experiment, we evaluated the capacity of *PhenoGMM* and *PhenoGrid* to both estimate and predict *α*-diversity of *in silico* synthetic microbial communities. A GMM of *K* = 128 or a fixed binning grid of dimensions 3 *×* 128 × 128 were fitted to the data. We conclude that *α*-diversity could be estimated properly, as predictions were significantly correlated with the true values according to Kendall’s *τ*_*B*_. *PhenoGMM* allowed better unsupervised *α*-diversity estimations compared to *PhenoGrid*, for which estimations were just above the significance level (*P* = 0.05) (Fig. 2A-C). Both approaches resulted either in comparable supervised predictions according to *τ*_*B*_ (Fig. 2D-F), or slightly in favor of *PhenoGMM* according to *R*^2^ (SI Fig. 1). Supervised predictions were considerably better than unsupervised estimations of *α*-diversity.

**Fig 2.**
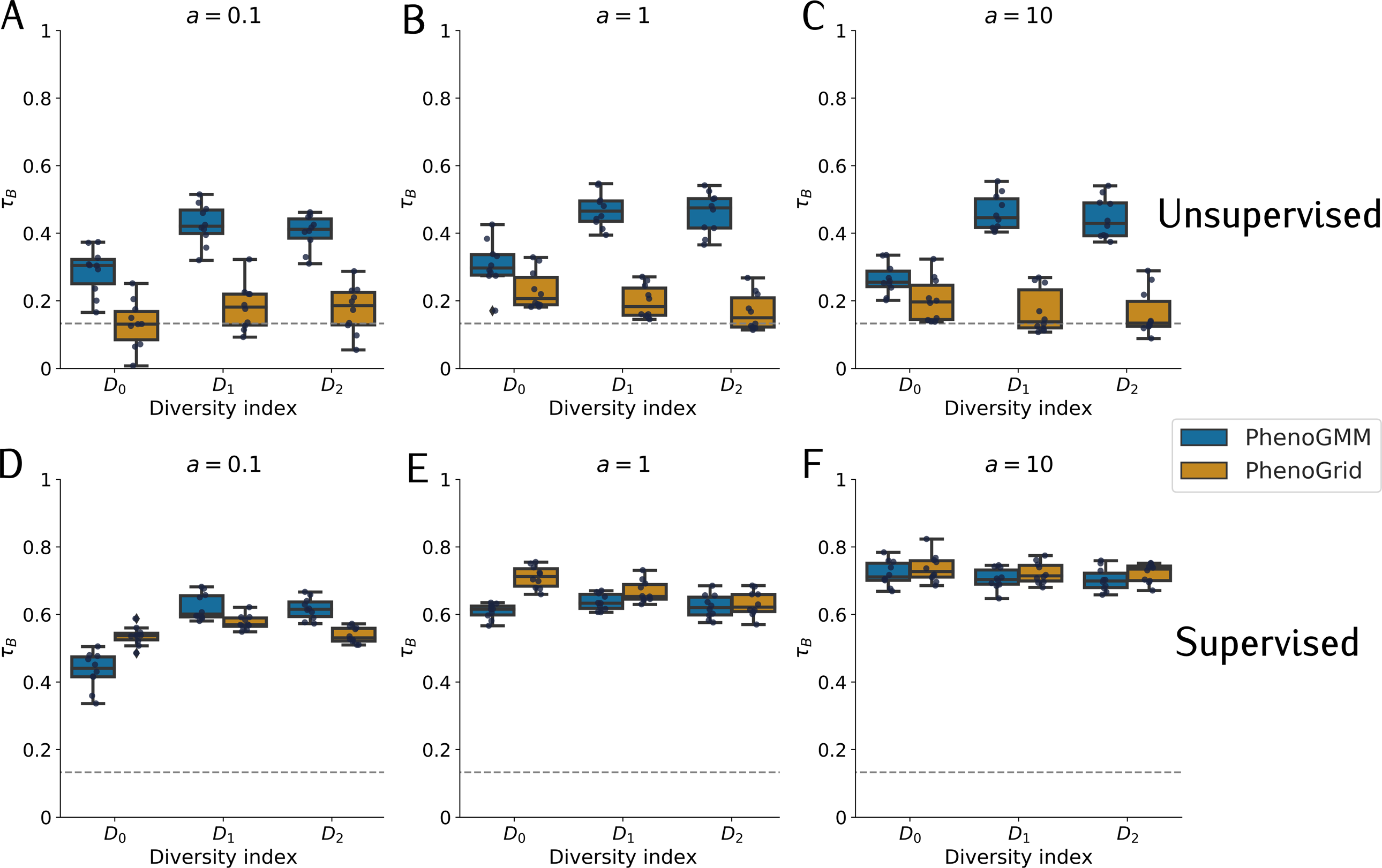
Summary of unsupervised and supervised *α*-diversity estimations for *in silico* synthetic microbial communities, quantified by Kendall’s *τ*_*B*_, for *PhenoGMM* and *PhenoGrid*. Both workflows were run ten times. Kendall’s *τ*_*B*_ was calculated between true and estimated values. Each boxplot displays the 25% and 75% quartiles of the *τ*_*B*_, and the whiskers show the full range of *τ*_*B*_. Each dot represents the resulting value from an individual run. **A-C**: Unsupervised estimations of *α*-diversity. **D-F**: Supervised estimations of *α*-diversity. **A**,**D**: *a* = 0.1; **B**,**E**: *a* = 1; **C**,**F**: *a* = 10. The dotted line indicates the strength of *τ*_*B*_ at *P* = 0.05.

We repeated the experiment for three different values of *a*, in which *a* determined how evenly the weight was spread amongst the different populations. If *a* was small, only a few species were dominantly present, if *a* was large, chances were higher that the weight was evenly spread amongst the different populations. This is illustrated using Lorenz curves, which depict the cumulative proportion of abundance versus the cumulative proportion of bacterial species (SI Fig. 2). We note that the predictive performance mainly depended on *a* and the diversity metric of choice. For example, the hardest setting was the one in which *a* = 0.1 and *D*_0_ the target variable to predict. In this case only a few populations made up a large part of the community (low *a*), but an equal weight was attributed to all species when defining diversity. Generally, if the abundance of populations was taken into account (*q >* 0), *α*-diversity predictions were better. In other words, FCM was able to capture community structure rather than the identity of the community.

We timed different steps in the workflow of *PhenoGMM* for *a* = 1 and *D*_1_. As there were 300 samples in our training set, this amounted to fitting a GMM to 1,5 million cells. The time in seconds was determined in function of the number of mixtures *K* (SI Fig. 3). Most importantly, the entire analysis remained under one hour. Most of the time was spent on fitting the GMM. Training a Random Forest model on the fitted GMM resulted in an average increase of 24,4% of the runtime for *K* = 256.

**Fig 3.**
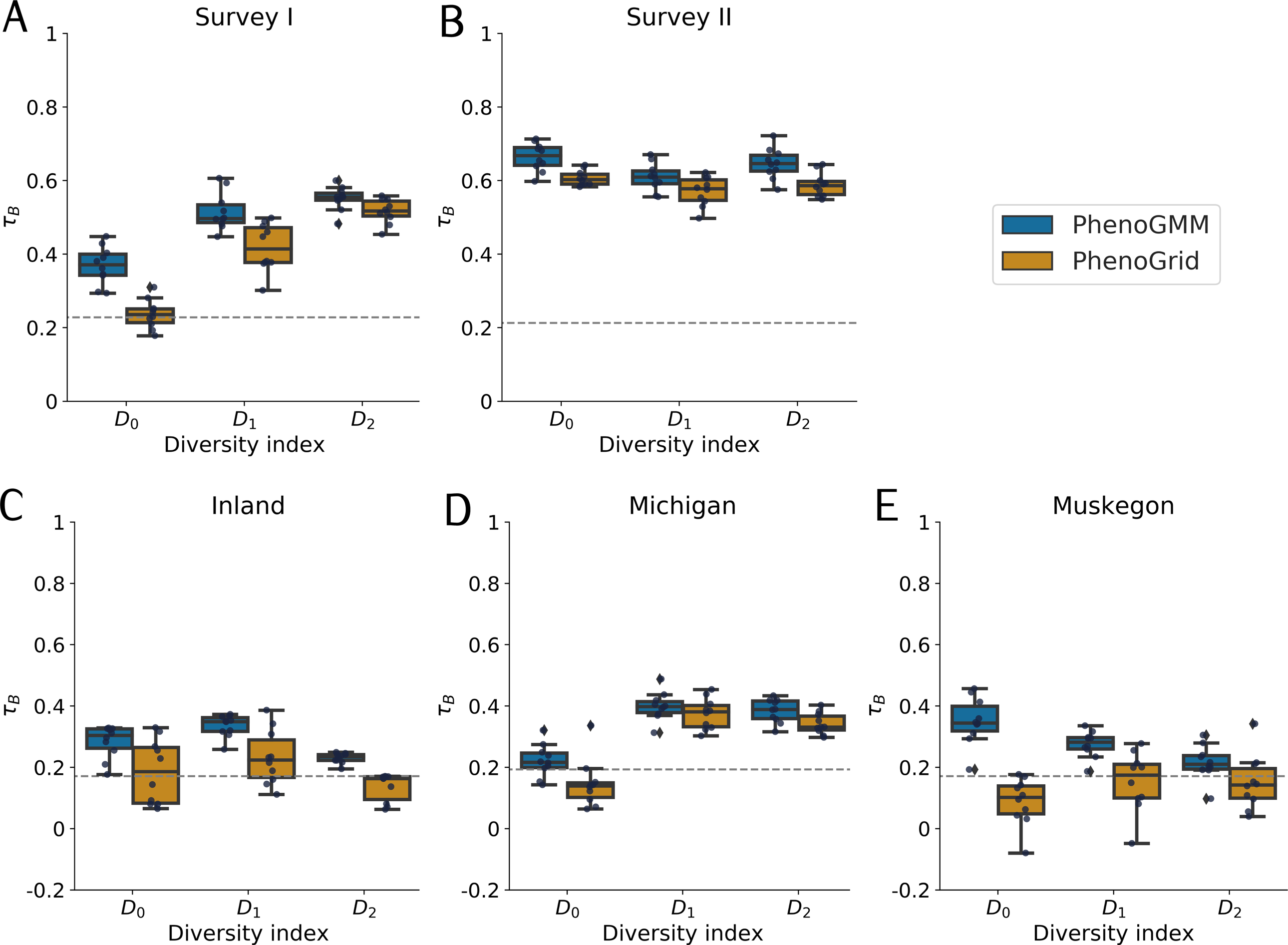
Summary of supervised *α*-diversity predictions for Dataset 2 and 3, evaluated by Kendall’s *τ*_*B*_, for *PhenoGMM* and *PhenoGrid*. Both methods were run ten times. Kendall’s *τ*_*B*_ was calculated between true and predicted diversity values. Each boxplot displays the 25% and 75% quartiles of the *τ*_*B*_, and the whiskers show the full range of *τ*_*B*_. Each dot represents the resulting value from an individual run. **A-B**: Results for Dataset 2; **A**: Survey I and **B**: Survey II. **C-E**: Results for the freshwater lake system microbiome. **C**: Inland lakes, **D**: Lake Michigan and **E**: Muskegon Lake. The dotted line indicates the strength of *τ*_*B*_ at *P* = 0.05.

In order to provide guidance concerning use of the model, the most important parameters were varied one by one (i.e., the number of included detectors *D*, the number of mixtures *K*, the number of cells sampled per file to fit a GMM denoted as N CELLS MIN, the number of cells sampled per individual sample to determine the cell counts per cluster denoted as N CELLS REP, a learning curve in function of N SAMPLES and the TYPE of covariance matrix used to fit a GMM). The performance was quantified using *R*^2^(*D*_1_) for *a* = 1 for a supervised analysis (SI Fig. 3). The results indicate that considering the predictive performance:

- *D*: including additional detectors improves the performance.
- *K*: generally, the higher *K*, the better the performance, which saturates after a specific threshold.
- N_CELLS_MIN: predictions are quite robust for this parameter.
- N_CELLS_REP: predictions are quite robust for this parameter.
- N_SAMPLES: predictive performance did not saturate yet at N SAMPLES = 300.
- TYPE: predictions are quite robust for the type of covariance matrix, but the ‘full’ type resulted in the best predictions.

### *PhenoGMM* allows to predict *α*-diversity for freshwater microbial communities

In the second experiment, we evaluated whether and to what extent it was possible to predict *α*-diversity for natural freshwater microbial communities, which were either part of a cooling water system (i.e. Survey I and II), or a freshwater lake system (i.e. Inland, Michigan and Muskegon). *α*-diversity values, based on 16S rRNA gene amplicon sequencing, were used as target variables to predict. Diversity predictions were feasible for Dataset 2 and in most cases for Dataset 3, i.e., significant according to *τ*_*B*_ (Fig. 3) and considerably higher than zero for *R*^2^(SI Fig. 5), but mostly depending on the dataset. It was easier to predict *α*-diversity for the cooling water microbial communities compared to the lake system microbial communities. In addition, predictions also depended on the diversity index. For example, it was easier to predict *D*_0_ compared to *D*_1_ or *D*_2_ for the Inland lakes. The predictive performance of *PhenoGMM* (*K* = 256) was better or similar compared to *PhenoGrid* (*K* = 3 *×* 128 *×* 128).

**Fig 4.**
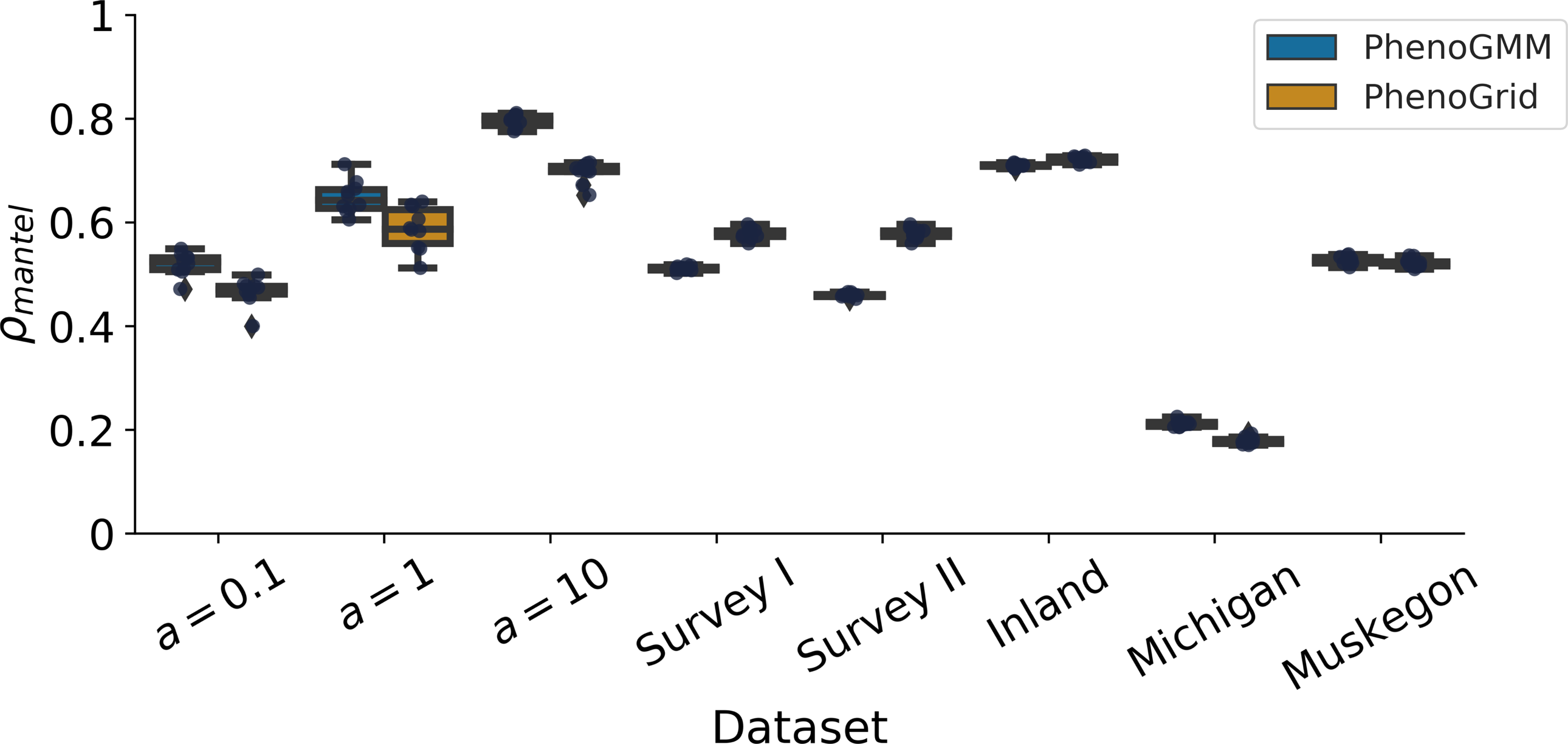
Summary of *β*-diversity estimations for Dataset 1, 2 and 3, evaluated by *ρ*_mantel_, for both *PhenoGMM* and *PhenoGrid*. Both methods were run ten times. *ρ*_mantel_ was calculated between the Bray-Curtis dissimilarity matrices based on cytometric fingerprints and the *in silico* community composition assembly strategy (Dataset 1) or 16S rRNA gene amplicon sequencing (Dataset 2 and 3). Each boxplot displays the 25% and 75% quartiles of *ρ*_mantel_, and the whiskers show the full range of *ρ*_mantel_.

**Fig 5.**
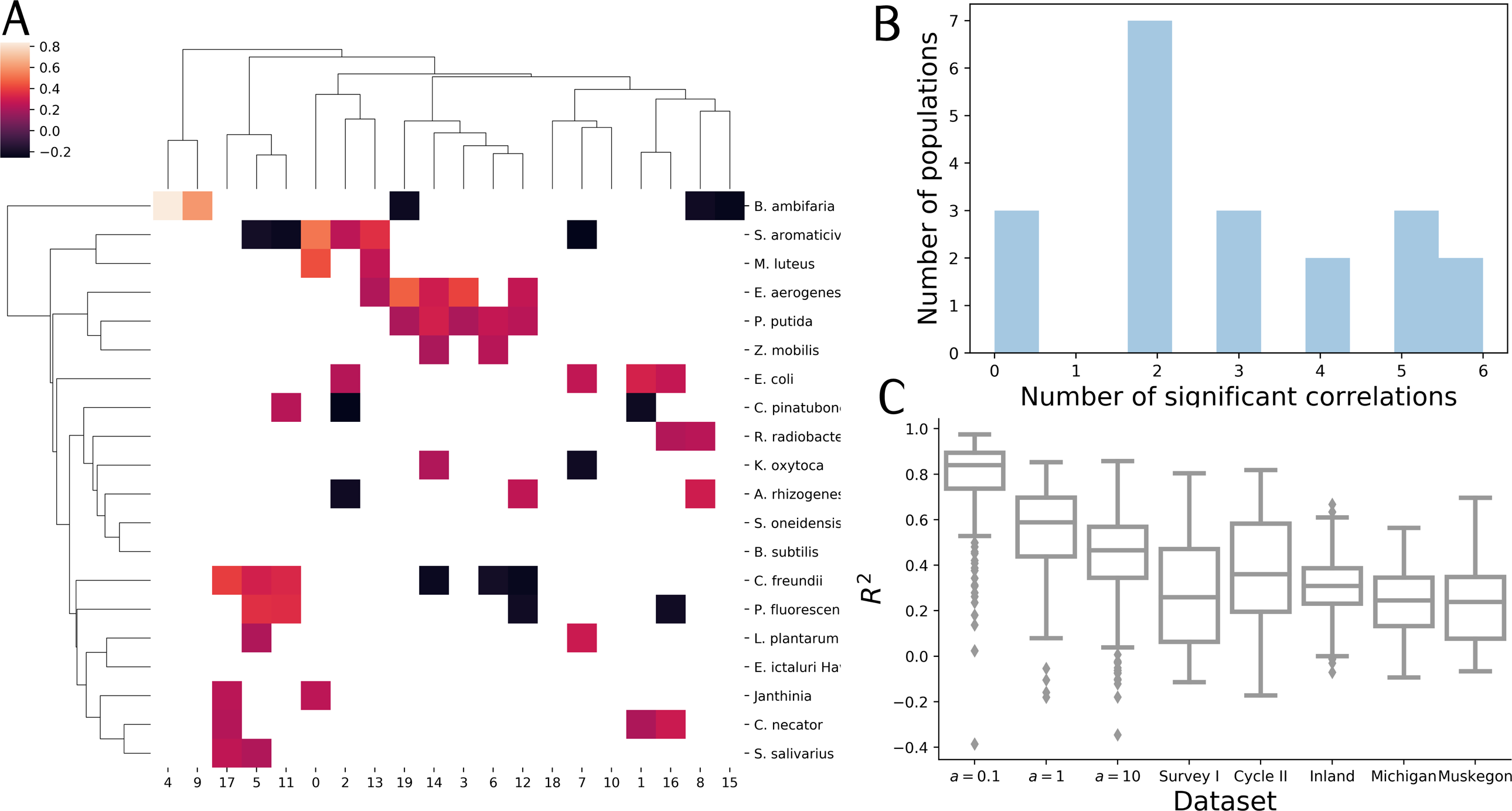
Summary of *PhenoGMM* abilities to predict OTU-abundances. **A:** Correspondence between variations in cell counts per mixture (columns) and abundances of bacterial populations (columns), quantified using the Kendall’s *τ*_*B*_. Values are given if *P ≤* 0.05, after performing a Benjamini-Hochberg correction for multiple hypothesis testing. **B:** Distribution of the number significant correlation with every mixture for each bacterial population. **C:** Predictions of taxon abundances for different datasets, expressed in terms of the *R*^2^ for the in silico datasets, based on a held-out test set, for the freshwater datasets the *R*^2^ OOB is reported. Boxplots represent the distribution of ten runs for 20 taxa. Each boxplot displays the 25% and 75% quartiles of *R*^2^, and the whiskers show the full range of *R*^2^, except for outliers in function of the interquartile range.

Unsupervised diversity estimations were evaluated as well (SI Fig. 6). Diversity estimations were highly significant for the cooling water microbiome, but were insignificant for two of the three freshwater lake systems according to both approaches (Inland and Lake Michigan). The diversity of the communities in Lake Muskegon could be successfully retrieved. If so, *PhenoGrid* outperformed *PhenoGMM* in most cases, indicating that even more mixtures might be needed to make it competitive with *PhenoGrid* in this setting. We conclude that FCM shows a moderate to strong connection with 16S rRNA gene sequencing data, depending on the dataset, which can be exploited using a supervised machine learning model.

### *PhenoGMM* allows estimations of *β*-diversity

We also evaluated the possibility to quantify *β*-diversity using *PhenoGMM*. This was done by calculating Bray-Curtis dissimilarities between the cytometric fingerprints and the OTU-tables. A mantel test was used to calculate the correlation between the cytometric dissimilarity matrix and the one derived from the *in silico* community composition (Dataset 1) or 16S rRNA gene sequencing (Datasets 2 and 3), for both *PhenoGMM* and *PhenoGrid* (Fig. 4). Both approaches resulted in statistically significant correlations (Mantel test, *P <* 0.05 for Lake Michigan, *P <* 0.001 for all other Datasets). *PhenoGMM* resulted in better estimations for Dataset 1, Lake Michigan and Muskegon Lake in Dataset 3, *PhenoGrid* was better for Dataset 2 and the Inland lakes in Dataset 3.

### *PhenoGMM* allows to predict individual bacterial abundances

The fact that microbial diversity can be estimated from cytometric data implies that the taxonomic structure of a microbial community is captured, at least to some extent, by the cytometric fingerprint. This opens up the opportunity to predict variations in abundance of individual bacterial populations as well. First, we constructed a fingerprint using 20 mixtures for Dataset 1 (*a* = 1) and correlated the relative cell counts per mixture with variations in individual abundances of bacterial populations (Fig. 5A). In most cases multiple clusters were correlated with multiple populations, which was due to the fact that bacterial populations exhibit overlapping cytometric fingerprints (Fig. 5B). At the same time, no cluster was correlated with all bacterial populations, motivating that despite the overlapping structure in a cytometric fingerprint, variations in the clusters could be related to variations in individual populations as well. The same procedure was applied for the Muskegon dataset, in which counts in 128 mixtures were correlated with the first 128 OTUs in the abundance table (SI Fig. 7). The same results were retrieved, meaning that almost every OTU showed a significant correspondence with cell count variations in multiple, but never all clusters.

Therefore, we tested whether we could predict the abundance of individual bacterial populations for all datasets (Fig. 5C). For Dataset 1, the individual abundances were known due to the experimental setup, for Dataset 2 and 3 we tested whether we could predict abundance values for the first 20 taxa in the OTU-table based on 16S rRNA gene amplicon sequencing data^2^. Predictions of taxon abundances were quantified in terms of the *R*^2^ on the test set for *in silico* synthetic communities or the out-of-bag error for natural communities, *R*^2^ OOB (Fig. 5C). Individual taxon abundances could be predicted based on cytometry data. For Dataset 1 this was possible for 81-100% of the populations, with *a* = 0.1 being the easiest setup to do so and *a* = 10 being the hardest setup to do so. This can be explained, as the weights of a composition will be divided over few species for *a* = 0.1, compared to compositions for *a* = 10. In other words, differences between individual abundances will be larger for *a* = 0.1, making it easier to predict them. For natural communities, we note that it was possible to capture and predict taxon abundances between 68.3-97.5% of the 20 most abundant taxa^3^.

## Discussion

In this paper we propose a more advanced cytometric fingerprinting strategy based on Gaussian Mixture Models (GMMs), which we called *PhenoGMM*. Our approach allows to create meaningful variables for microbial cytometry data in order to describe community dynamics and reduces the number of -describing variables considerably compared to traditional binning approaches. This makes the use of predictive models, in this study by means of Random Forest regression, more feasible. We evaluated the performance of *PhenoGMM* both for unsupervised estimations and supervised predictions of microbial diversity using multiple datasets stemming from synthetic and freshwater microbial communities. We compared it with the performance of a generic traditional binning approach, which we for this work called *PhenoGrid*.

In the first part of the paper, we constructed communities *in silico* by aggregating cytometric characterizations of individual bacterial populations according to different predefined compositions. This allowed us to simulate microbial community compositions in a highly precise and controlled way. Upon making predictions, *PhenoGMM* resulted in either more or equally accurate predictions compared to *PhenoGrid* for all settings. In other words, cytometric fingerprints allow to quantify the diversity of synthetic microbial communities. The total analysis time of *PhenoGMM* remains under one hour for the analysis of one million cells.

In the second part we showed that FCM data can be used to predict diversity values based on 16S rRNA gene sequencing data. Supervised predictions of *α*-diversity resulted in higher correlations with the target diversity values for *PhenoGMM* compared to *PhenoGrid*, while this was the other way around when performing an unsupervised analysis. Note that we do not expect to find a ‘perfect’ correlation with taxonomic diversity. Besides the fact that 16S rRNA gene amplicon sequencing is subject to a number of biases [35, 38], microbial FCM captures both taxonomic and physiological changes. Therefore, the strength of the correspondence between cytometric and taxonomic diversity will vary from system to system. The levels of trophicity differ between datastes, which could explain why the estimations and predictions for Dataset 2 (which originates from an oligotrophic environment, see [45]) are better than those for Dataset 3 (which originate samples from lakes of varying trophicity, see [53]). Another reason could be due to the complexity of the datasets. Dataset 2 is less complex, containing many measurements over a few days, compared to Dataset 3, which contains samples that span a much larger range in time (years) and space (multiple locations). Due to limitations in sample size, the predictive performance for these datasets was evaluated using the out-of-bag error of the best Random Forest model from a blocked cross-validation scheme. Future research is recommended to assess and confirm the correspondence between cytometric and taxonomic diversity in other ecosystems and by means of larger datasets, allowing the use of a separate test set.

We overcluster the data to model the multiple and highly overlapping cell distributions. However, this makes it difficult to determine the exact number of populations. As the number of mixtures *K* increases, the performance saturates gradually, and more mixtures will not improve predictions. *PhenoGMM* could also be tailored towards the identification of separated cell populations, for example to identify phytoplankton populations (conform [28]) or for the identification of so-called high and low nucleic acid groups (conform [21]). In this case, if the number of populations is known beforehand, *K* can be chosen accordingly; if this is not known, one can use decision rules such as the Bayesian Information Criterion (BIC) to determine the optimal number of mixtures (conform [36]).

Few reports exist that quantitatively evaluate fingerprinting approaches for the analysis of microbial cytometry data. A brief comparison study with *n* = 21 samples has been recently conducted [39], illustrating a better performance for *FlowFP* [50], compared to the use of *FlowCyBar* [30]. *FlowFP* is quite similar compared to *PhenoGMM*, as it makes use of an adaptive binning approach, in which bins are smaller when the density of the data is higher, while *FlowCyBar* makes use of manually annotated clusters. However, the bins are still rectangular in shape, while *PhenoGMM* allows clusters to be of any shape. Most fingerprinting strategies make use of manual annotation of clusters or of fixed binning approaches (see e.g. the report by [31] which qualitatively discusses different existing methods). In almost all cases, only bivariate interactions are inspected. *PhenoGMM* allows to model the full parameter space at once. This is interesting, because although it is hard to develop multicolor approaches for bacterial analyses, they are possible (see e.g. the work by [4]). In addition, our research group has established that additional detectors that capture signals due to spillover can assist in the discrimination between bacterial species [52]. Therefore, the parameter space in which bacterial cells can be described is increasing, and *PhenoGMM* is able to model this straightforwardly. Because it is in an adaptive strategy as well, by defining small clusters in regions of high density and vice versa, it reduces the number of sample-describing variables considerably compared to fixed binning approaches. Other adaptive binning strategies have been proposed for microbial FCM data as well, however these still only investigate bivariate interactions [3, 27].

Our approach comes with a number of caveats. First, *PhenoGMM* fits a fingerprint template based on the concatenation of measured samples. New samples are characterized based on this template. In case multiple samples diverge considerably from those that were used to determine the template (for example in case an experiment was conducted in different conditions), we recommend to refit the model. Second, *PhenoGMM* overclusters the data, which might result in a number of correlated variables. We recommend therefore researchers to use a classification or regression method that is able to deal with multicollinearity, which is why we used Random Forest regression in this work. Other methods that might be suitable are regularized regression methods, such as the Lasso or ElasticNet [57, 64]. Third, although the performance tends to saturate once *K* is high enough, this threshold seems to be application dependent, and one needs to validate the settings of the approach.

Our *in silico* benchmark study made use of cytometric characterizations of individual bacterial populations. These populations are known to exhibit considerable heterogeneity due to cell size diversity and cell cycle variations [61]. Our research group has recently shown that the cytometric diversity of an individual population reduces when that population is part of a co-culture [25]. Therefore, data used for the *in silico* community creation setup cannot be used to study environmental samples, as we hypothesize that members of natural communities will have a different cytometric fingerprint as opposed to populations that were grown and measured individually. Yet we believe that our *in silico* approach is useful, as it allows to simulate variations in cytometric community structure with high precision.

In this study we focused mainly on estimations of *α*-diversity (i.e., within-sample diversity), but quantification of *β*-diversity (i.e. between-sample diversity) can be successfully performed as well. In addition, it is possible to predict variations in the abundance of a specific bacterial populations. This might be interesting for certain biotechnological applications, in which researchers or engineers are not interested in the total diversity of the community but in the behavior of a specific bacterial population.

Technological advancements have enabled an automation of the data acquisition, resulting in a detailed characterization of the microbial community on-line (i.e., samples are measured at routine intervals between 5-15 min) or even in real-time (i.e., near-continuous measurements) [6, 24]. Therefore we see great potential to use FCM as a monitoring technique to rapidly and frequently investigate microbial community dynamics. In this work we have confirmed the strong correspondence between FCM and the genetic make-up of a community, quantified by 16S rRNA gene sequencing. Machine learning models can be used to exploit the relationship between the two. However, they can also be used to perform classification at the community level, to for example categorize communities according to the system they are a part of [16, 18], or to identify a case versus control status.

To conclude, *PhenoGMM* allows to extract meaningful information from microbial FCM measurements. Besides the fact that it facilitates the use of supervised machine learning models, cytometric diversity estimations can also be assessed directly. It has to be noted that quantification of diversity should serve as a starting point to test ecological hypotheses rather than as a final outcome of an experiment [55]. Microbial FCM, in combination with *PhenoGMM*, has the potential to be an effective strategy to serve this research line in microbial ecology.

## Acknowledgments

The computational resources (Stevin Supercomputer Infrastructure) and services used in this work were provided by the VSC (Flemish Supercomputer Center), funded by Ghent University, the Hercules Foundation and the Flemish Government department EWI.

## Supporting information

**S1 Fig. Summary of supervised** *α***-diversity estimations for *in silico* synthetic microbial communities, quantified by** *R*^2^, **using *PhenoGMM* and *PhenoGrid***. *PhenoGMM* and *PhenoGrid* were run ten times. The *R*^2^ was calculated between true and estimated values. Each boxplot displays the 25% and 75% quartiles of the *τ*_*B*_, and the whiskers show the full range of *R*^2^. **A**: *a* = 0.1, **B**: *a* = 1 and **C**: *a* = 10.

**S2 Fig.Lorenz curves for all sampled *in silico* communities for a** = **0.1, 1, 10**. A: training set (300 communities), B: test set (100 communities)

**S3 Fig. Benchmarking of *PhenoGMM* in function of the time in seconds**. Each analysis was run on a separate node of a computer infrastructure, with 2.6 Ghz CPU and 20GB of RAM for each node. **A**: Time to fit a GMM model and perform unsupervised diversity estimations according to equation 3 (blue line), evaluated by *τ*_*B*_(*D*_1_) (orange line). **B**: Time to fit a GMM model, fit a Random Forest regression model and perform supervised diversity predictions of *D*_1_ (blue line), evaluated by *τ*_*B*_(*D*_1_) (orange line). *PhenoGMM* was run five times for different *K*, for which the mean and corresponding standard deviation are visualized.

**S4 Fig.Influence of different parameters of *PhenoGMM* on both supervised and unsupervised estimations of** *D*_1_ **for** *a* = 1, **quantified by Kendall’s** *τ*_*B*_. Declaration of parameters: *D*: number of included detectors (height-signal); *K*: number of mixtures; N Cells c: number of cells that are sampled per community and concatenated (c) to fit a GMM; N Cells i: number of cells that are sampled per individual (i) community to derive cell counts using a fitted GMM; TYPE: type of GMM that is used.

**S5 Fig.Summary of supervised** *α***-diversity out-of-bag predictions for Dataset 2 and 3, evaluated by the** *R*^2^, **using both *PhenoGMM* and *PhenoGrid***. Both methods were run ten times. The *R*^2^ was calculated between true and out-of-bag predicted diversity values. Each boxplot displays the 25% and 75% quartiles of the *R*^2^, and the whiskers show the full range of the *R*^2^. **A-B**: Results for Dataset 2; **A**: Survey I and **B**: Survey II. **C-E**: Results for the freshwater lake system microbiome. **C**: Inland lakes, **D**: Lake Michigan and **E**: Muskegon Lake.

**S6 Fig.Summary of unsupervised** *α***-diversity estimations for Dataset 2 and 3, evaluated by Kendall’s** *τ*_*B*_, **using both *PhenoGMM* and *PhenoGrid***. Both methods were run ten times. Kendall’s *τ*_*B*_ was calculated between true and estimated diversity values. Each boxplot displays the 25% and 75% quartiles of the *τ*_*B*_, and the whiskers show the full range of *τ*_*B*_. **A-B**: Results for Dataset 2; **A**: Survey I and **B**: Survey II. **C-E**: Results for the freshwater lake system microbiome. **C**: Inland lakes, **D**: Lake Michigan and **E**: Muskegon Lake. The dotted line indicates the strength of *τ*_*B*_ at *P* = 0.05.

**S7 Fig.Summary of the correspondence between cytometric fingerprints and OTU-abundances for the Muskegon dataset. A:** Correspondence between variations in cell counts per mixture (columns) and abundances of bacterial populations (columns), quantified using the Kendall’s *τ*_*B*_. Values are given if *P ≤* 0.05, after performing a Benjamini-Hochberg correction for multiple hypothesis testing. **B:** Distribution of the number significant correlations with every mixture for each bacterial population.

## Footnotes

1 Note, this is not the same as *k*-means clustering. In this case, all mixtures would share the same single variance.

2 Except for Survey I, for which results are presented for 18 taxa due to the fact that two taxa did not vary in abundance, and therefore resulted in ‘perfect’ predictions

3 These numbers were calculated by counting the number of OTUs for all runs that resulted in a positive and significant *τ*_*B*_ (*P <* 0.05) after performing a Benjamini-Hochberg correction for multiple hypothesis testing.

