## Supplementary Figures for "*PhenoGMM*: Gaussian mixture modelling of microbial cytometry data enables efficient predictions of biodiversity"

### Supporting Figures

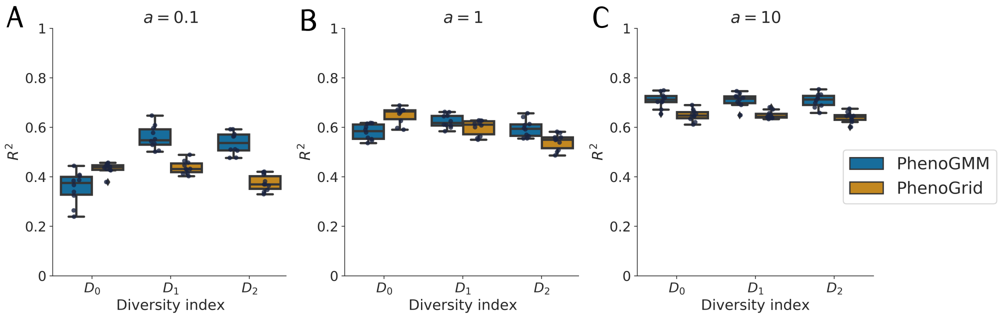


**Fig 1. Summary of supervised $\alpha$-diversity estimations for *in silico* synthetic microbial communities, quantified by $R^2$, using *PhenoGMM* and *PhenoGrid*.** *PhenoGMM* and *PhenoGrid* were run ten times. The $R^2$ was calculated between true and estimated values. Each boxplot displays the 25% and 75% quartiles of the $\tau_B$, and the whiskers show the full range of $R^2$. Each dot represents the resulting value from an individual run. **A**: $a = 0.1$, **B**: $a = 1$ and **C**: $a = 10$.

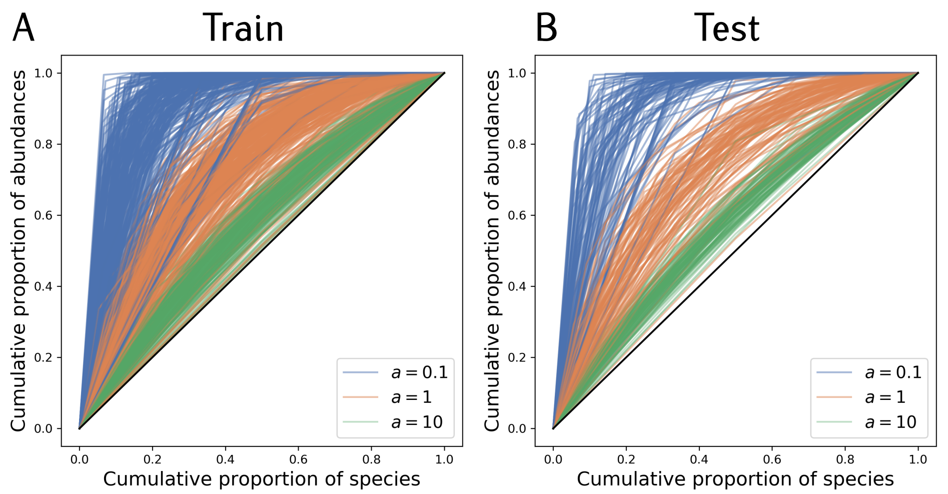


**Fig 2. Lorenz curves for all sampled *in silico* communities for $\mathbf{a = 0.1, 1, 10}$.** A: training set (300 communities), B: test set (100 communities).

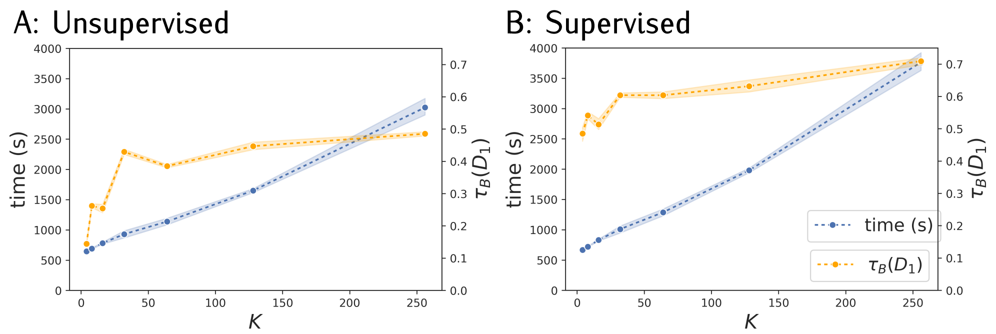


**Fig 3. Benchmarking of *PhenoGMM* in function of the time in seconds.** Each analysis was run on a separate node of a computer infrastructure, with 2.6 Ghz CPU and 20GB of RAM for each node. **A**: Time to fit a GMM model and perform unsupervised diversity estimations according to equation 3 (blue line), evaluated by $\tau_B(D_1)$ (orange line). **B**: Time to fit a GMM model, fit a Random Forest regression model and perform supervised diversity predictions of $D_1$ (blue line), evaluated by $\tau_B(D_1)$ (orange line). *PhenoGMM* was run five times for different $K$, for which the mean and corresponding standard deviation are visualized.

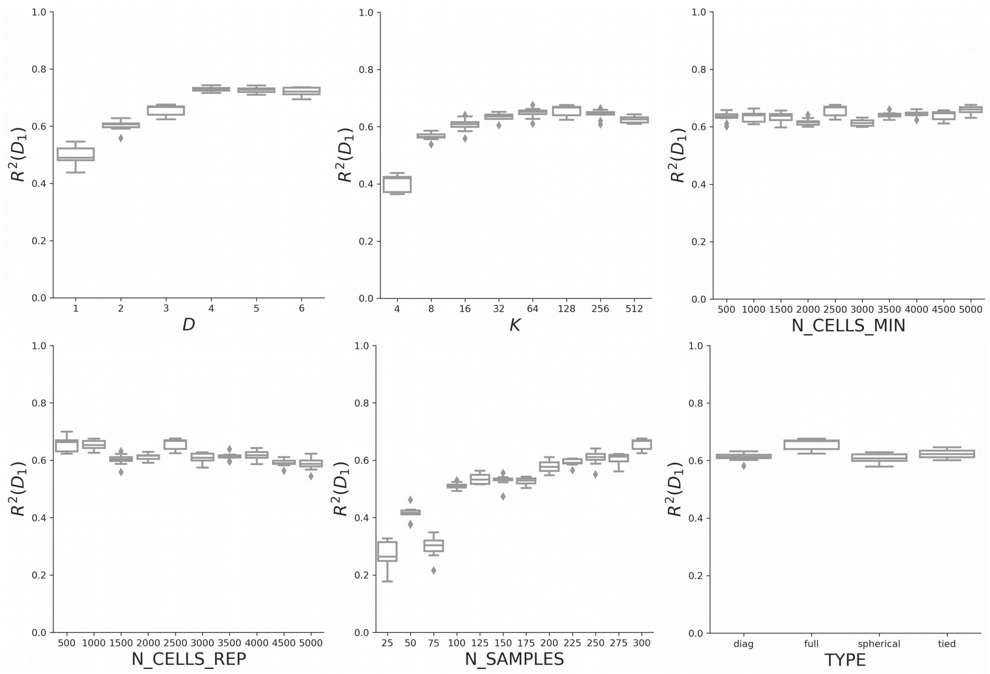


**Fig 4. Influence of different parameters of *PhenoGMM* on both supervised and unsupervised estimations of $D_1$ for $a = 1$, quantified by Kendall's $\tau_B$.** Declaration of parameters: $D$: number of included detectors (height-signal); $K$: number of mixtures; N_Cells_c: number of cells that are sampled per community and concatenated (c) to fit a GMM; N_Cells_i: number of cells that are sampled per individual (i) community to derive cell counts using a fitted GMM; TYPE: type of GMM that is used.

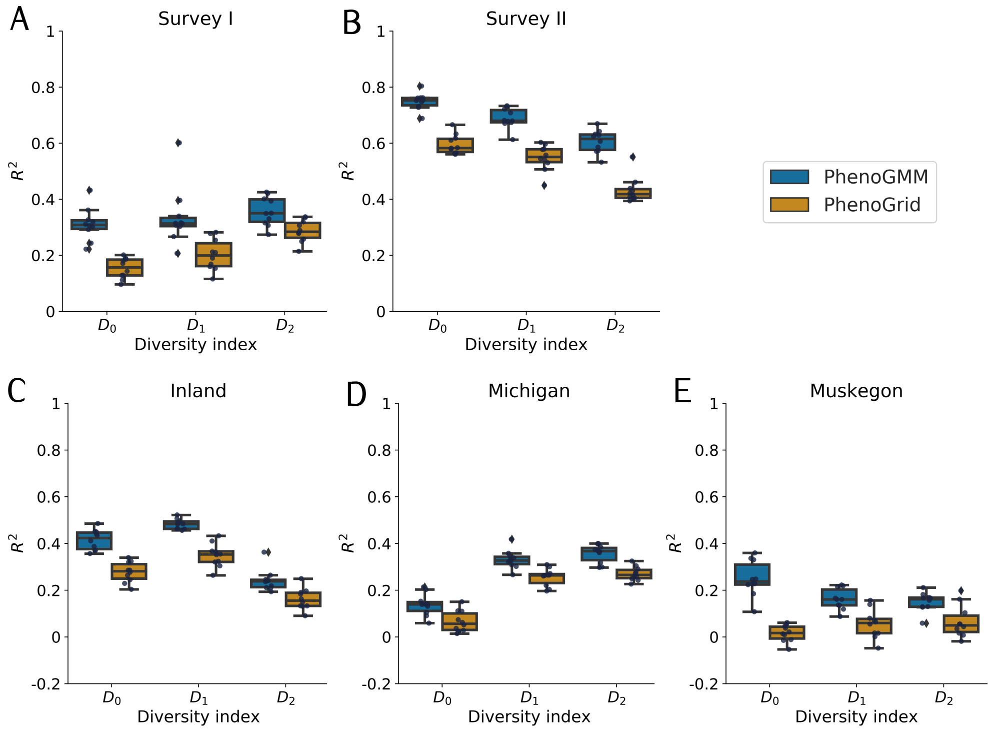


**Fig 5. Summary of supervised $\alpha$-diversity out-of-bag predictions for Dataset 2 and 3, evaluated by the $R^2$, using both *PhenoGMM* and *PhenoGrid*.** Both methods were run ten times. The $R^2$ was calculated between true and out-of-bag predicted diversity values. Each boxplot displays the 25% and 75% quartiles of the $R^2$, and the whiskers show the full range of the $R^2$. Each dot represents the resulting value from an individual run. **A-B**: Results for Dataset 2; **A**: Survey I and **B**: Survey II. **C-E**: Results for the freshwater lake system microbiome. **C**: Inland lakes, **D**: Lake Michigan and **E**: Muskegon Lake.

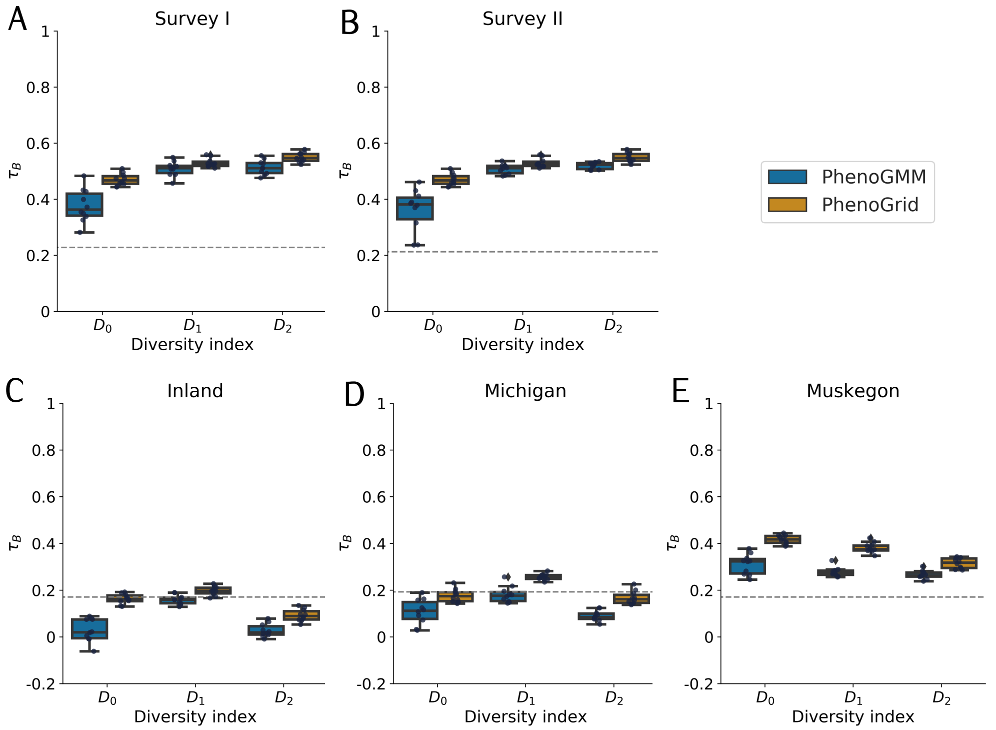


**Fig 6. Summary of unsupervised $\alpha$-diversity estimations for Dataset 2 and 3, evaluated by Kendall's $\tau_B$, using both *PhenoGMM* and *PhenoGrid*.** Both methods were run ten times. Kendall's $\tau_B$ was calculated between true and estimated diversity values. Each boxplot displays the 25% and 75% quartiles of the $\tau_B$, and the whiskers show the full range of $\tau_B$. Each dot represents the resulting value from an individual run. **A-B**: Results for Dataset 2; **A**: Survey I and **B**: Survey II. **C-E**: Results for the freshwater lake system microbiome. **C**: Inland lakes, **D**: Lake Michigan and **E**: Muskegon Lake. The dotted line indicates the strength of $\tau_B$ at $P = 0.05$.

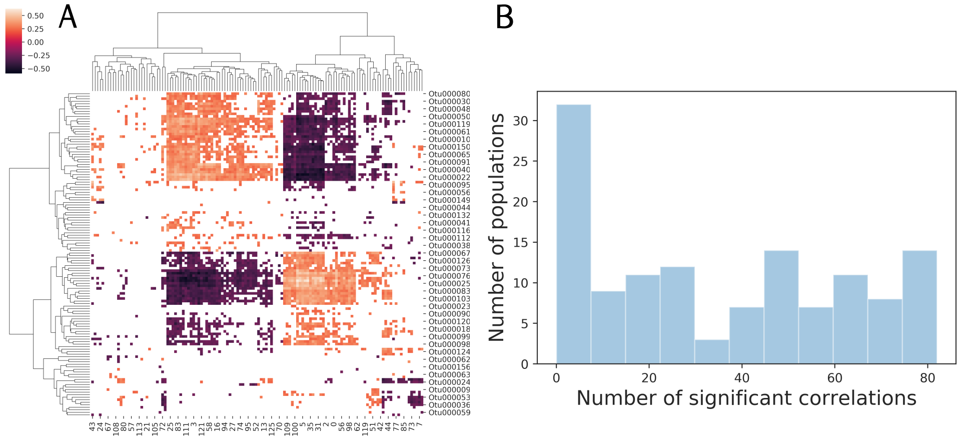


**Fig 7. Summary of the correspondence between cytometric fingerprints and OTU-abundances for the Muskegon dataset**. **A:** Correspondence between variations in cell counts per mixture (columns) and abundances of bacterial populations (columns), quantified using the Kendall's $\tau_B$. Values are given if $P \leq 0.05$, after performing a Benjamini-Hochberg correction for multiple hypothesis testing. **B:** Distribution of the number significant correlations with every mixture for each bacterial population.
